# Comparative Performance of Scribe and Database Search Engines in Metaproteomic Profiling of a Ground-Truth Microbiome Dataset

**DOI:** 10.1101/2025.05.15.654320

**Authors:** Andrew T. Rajczewski, Subina Mehta, Reid Wagner, Wassim Gabriel, James Johnson, Katherine Do, Simina Vintila, Mathias Wilhelm, Manuel Kleiner, Brian C. Searle, Timothy J. Griffin, Pratik D. Jagtap

## Abstract

Mass spectrometry-based metaproteomics, the identification and quantification of thousands of proteins expressed by complex microbial communities, has become pivotal for unraveling functional interactions within microbiomes. However, metaproteomics data analysis encounters many challenges, including the search of tandem mass spectra against a protein sequence database using proteomics database search algorithms. We used a ground-truth dataset to assess a spectral library searching method against established database searching approaches. Mass spectrometry data collected by data-dependent acquisition (DDA-MS) was analyzed using database searching approaches (MaxQuant and FragPipe), as well as using Scribe with Prosit predicted spectral libraries. We used FASTA databases that included protein sequences from microbial species present in the ground-truth dataset along with background protein sequences, to estimate error rates and assess the effects on detection, peptide-spectral match quality, and quantification. Using the Scribe search engine resulted in more proteins detected at a 1% false discovery rate (FDR) compared to MaxQuant or FragPipe, while FragPipe detected more peptides verified by PepQuery. Scribe was able to detect more low-abundance proteins in the microbiome dataset and was more accurate in quantifying the microbial community composition. This research provides insights and guidance for metaproteomics researchers aiming to optimize results in their analysis of DDA-MS data.

## INTRODUCTION

Metaproteomics aims to characterize the protein complement of a microbial community, providing insights into both the taxonomic composition and functional roles of the microbes within that community^1,2^. This characterization is crucial for understanding the complex interactions and dynamics of microbial ecosystems in various contexts, such as the human microbiome or environmental samples.

Researchers typically rely on tandem mass spectrometry (MS/MS) data to identify and quantify proteins present in a sample in metaproteomics analyses^3–5^. MS/MS data can be acquired via either a) data-dependent acquisition (DDA-MS), which selects and fragments the most intense peptide ions, or b) data-independent acquisition (DIA-MS), which fragments all ions within predefined m/z windows. While the potential benefits of a DIA-MS approach have been highlighted^6–8^, most metaproteomics researchers currently use a DDA-MS approach. Shotgun proteomics involves analyzing a complex mixture of proteins by breaking them into peptides through proteolysis, separating the mixtures by liquid chromatography (LC), and characterizing these peptides with tandem mass spectrometry, and inferring parent proteins by matching tandem mass spectra to a reference database.

Two principal strategies exist for matching spectra to peptide sequences. The conventional approach to matching spectra to peptides involves searching acquired DDA-MS-generated MS/MS spectra against theoretical spectra generated from peptide sequences derived from a FASTA protein sequence database^9,10^. The protein sequence database is ideally constructed from genome sequences of the organisms in the sample^11^, and the search identifies the microbial peptides present in the sample. While several algorithms exist for matching the empirical and theoretically-derived tandem mass spectra in database searches^10^, it should be noted that the theoretical spectra lack information concerning the intensities of the product ions and include all possible product ions; these factors introduce the potential for the misidentification of peptides^12^. An alternate strategy for identifying spectral matches is spectral library searching^12,13^, wherein the acquired MS/MS spectra are searched against and matched to a library of confidently annotated peptide spectra observed in previous experiments. These library spectra reflect the relative abundances of the product ions in each peptide spectra as well as the presence of non-standard ions such as water and ammonia losses^14^ not predicted in the theoretical spectra of sequence database searching, allowing for the potential for more high-confidence matching of spectra^15^.

Despite its promise as a research tool, metaproteomics faces unique challenges, particularly when dealing with highly diverse microbiomes^16,17^. If the used protein sequence database is very large, this can lead to reduced sensitivity in matching spectra to peptides, hindering the accurate identification of proteins. Although iterative search strategies have been utilized^18–22^, the risk of false positives, where MS/MS spectra are incorrectly matched to peptides, becomes more pronounced with a larger and more diverse database^23,24^. We hypothesize that a spectral library searching strategy would mitigate the problem of false positives due to the increased sensitivity of this method associated with the ability to assign peptides to low-quality and complex spectra^15^. A limit of spectral library searching has historically been a lack of high-quality spectra with which to construct libraries; this can be rectified with computationally predicted peptide fragmentation patterns to generate more comprehensive spectral libraries^25^. The use of deep learning technology has enabled the construction of spectral libraries complete with fragment ion intensity predictions and peptide retention times, allowing for even more confident identifications of peptides^26^. The ability to search MS data and quantify proteins has been leveraged in the new spectral library search engine Scribe by Searle *et al*.^27^

In this study, we conducted a comparative analysis between spectral library searching with Scribe and conventional peptide-spectrum matching using the commonly used DDA-MS-based pipelines MaxQuant^28^ and Fragpipe^29^. We explored challenges arising in metaproteomics searches, in particular in handling large and diverse microbiomes. Spectral library searching, exemplified by Scribe, offers a promising alternative, addressing these challenges and providing a more sensitive and accurate means of identifying proteins within complex microbial ecosystems.

## METHODS

### Synthetic Microbiome sample

A ground-truth dataset of a digested mixture of 30 microbial species (32 microbial strains; three *Salmonella enterica typhimurium* LT2 strains were considered as a single species LT2) of Archaea, Bacteria, Eukaryotes, and Bacteriophages with known species abundances was used for all analyses (UNEVEN community^30^; See **Supplementary Table 1**). Microbial communities were constituted in the Kleiner Lab at North Carolina State University for a previous study30. Briefly, 32 microbial cultures of archaea, bacteria, and phages were washed in phosphate-buffered saline pH = 7.4 to remove residual culture media and aliquoted to generate cell pellets. Replicates of each species’ cell pellet were lysed, and protein concentrations were determined. Cell pellets were reconstituted in ultrapure water and combined to generate multiple aliquots of each community before being snap-frozen. For protein isolation and digestion, Tris-HCl buffer pH = 7.6 supplemented with 4% SDS and 0.1M DTT was added to each frozen community sample in a 1:10 ratio before the samples were disrupted by bead beating. Samples were then incubated at 95°C for 10 minutes, after which cellular debris was removed by centrifugation at 21,000g for 5 minutes. Following lysis, 60 μL of cleared lysate was used for digestion using a modified filter-aided sample preparation (FASP) protocol described by Wisniewski *et al*.^31^ Lysate was combined with 400 μL of UA buffer (8 M urea in 0.1 M Tris-HCl, pH 8.5) twice and loaded onto 10 kDa molecular weight cutoff 500 μl centrifugal filters (VWR International) by centrifuging for 15-20 minutes at 14,000g. Filters were then washed with 200 μL of UA buffer and centrifuged for 15-20 minutes. IAA solution (0.05 M iodoacetamide in UA buffer, 100 μL) was added to the filters and incubated for 20 minutes prior to a 20-minute centrifugation. Filters were then washed 3 times with 100 μL of UA buffer, followed by another 3 washes with 100 μL of ABC buffer (50□mM ammonium bicarbonate). A digestion mixture of 1 μg MS-grade trypsin (Thermo Scientific Pierce, Rockford, IL) in 40□μl of ABC buffer was added to each filter and incubated overnight at 37°C. The following day, tryptic peptides were eluted using 50□μl of 0.5 M NaCl and 20 minutes of centrifugation. The resulting peptides were desalted using Sep-Pak C18 Plus Light Cartridges (Waters, Milford, MA, USA) in accordance with the manufacturer’s instructions. Peptide concentrations were determined using the Pierce Micro BCA assay (Thermo Scientific Pierce, Rockford, IL, USA) following the manufacturer’s instructions.

Based on protein sequence similarity between strains of the same species, 30 species were used for evaluation. This UNEVEN mock community of 30 microbial species was designed to cover a large range of species abundances, both at the level of cell number and proteinaceous biomass, to test for the dynamic range and detection limits of the quantification methods.

### LC-MS/MS conditions

Four replicates of the UNEVEN mock community were analyzed by LC-MS/MS. Water with 0.1% formic acid was used as mobile phase A and acetonitrile with 0.1% formic acid was mobile phase B for LC applications. Samples were analyzed on a QExactive Quadrupole Orbitrap Hybrid Mass Spectrometer interfaced with an Ultimate 3000 UHPLC run in nano mode and plumbed with a nanoLC column packed with Luna C18 5µm resin (15 cm x 75 µm). For all LC-MS runs the same gradient was used, wherein the relative composition was held constant at 5% B for 5 minutes, followed by an increase to 35% B from 5 minutes to 95 minutes, an increase to 95% B from 95 to 100 minutes, a constant composition of 95% B from 100 to 110 minutes, a decrease from 95% B to 5% from 110 to 112 minutes, and a final re-equilibration stage held at 5% B from 112 minutes to 120 minutes. The instrument was run in positive mode using Full MS/dd-MS^2^ Top 15 mode. For the Full MS scan, the resolution was 35,000 with an AGC target of 1e6, a maximum IT of 30 milliseconds, and a scan range of 400-1600 m/z. Data-dependent MS^2^ were collected at a resolution of 17,500 with an AGC target of 1e6, a maximum IT of 50 milliseconds, an isolation window of 2.0 m/z, and a scan range of 200 - 2000 m/z.

### Construction of protein sequence databases

A composite protein sequence database (112,580 protein sequences from 30 microbial species) was generated by combining the protein sequences of all species in the UNEVEN mock community into a single database. Protein sequences were acquired from UniProt or NCBI and are detailed in Supplement Table 1. The protein database was clustered at 95% identity to remove redundant sequences using cd-hit^32^. We have added the common Repository of Adventitious Proteins, a list of 116 proteins commonly found in proteomics experiments that are present through unavoidable contamination of protein samples. This protein sequence database was named 1X database (112,580 protein sequences) in Results and Figures. 112,580 protein sequences from the ‘background’ Integrated Gene Catalog (IGC) Database^33^ were randomly selected and merged with the 1X database and named the 2X database (225,160 protein sequences). The IGC contains a large collection of genes from human microbiomes.

### Construction of spectral library

For Scribe spectral library searches, the protein sequence databases (1X and 2X protein sequence FASTA files) were converted into predicted spectral libraries (.dlib format) using Oktoberfest^34^. The spectral library was generated using the Prosit 2020 Intensity HCD model^35^ with up to 2 allowed miss cleavages, collision energy 33, peptide length between 7 to 30, and up to 3 oxidized methionines were permitted.

### Search algorithms

Raw DDA-MS files were searched against the protein sequence databases 1X and 2X using either MaxQuant^28^ (version 2.6.7.0) or FragPipe^29^ (version 22). For spectral library searches, MSconvert generated mzML files of the Raw files were used for searches using Scribe^27^ (version 2.12.30). For all searches, the precursor tolerance was set to 20 ppm and the fragment tolerance was set to 15 ppm. For spectrum matching, b- and y-ions were selected. Methionine oxidation was set as a variable modification while carbamidomethylation of cysteine was set as a fixed modification. The three search algorithms were used for peptide and protein identification at 1% Global FDR from four replicates of DDA-MS datasets. In all three algorithms, a target-decoy search approach was used with a reverse decoy database to estimate the FDR based on Posterior Error Probability (PEP) scores (MaxQuant) or probability scores computed by PeptideProphet (FragPipe). Outputs from the search algorithms were used to report peptides and proteins from the microbial mix as well as the background IGC proteins. Deep Venn^36^ was used to generate the Venn diagrams reported. Moreover, BLASTP analysis was used to assign background peptides to the microbial mix proteins by searching them against the 1X database. The outputs from search algorithms were also used for protein and peptide quantification relative to the known abundances of each species when the microbiome was created.

### Peptide Verification

To assess the quality of the peptide-spectrum assignment, PepQuery software was used as an alternative approach to confirm the presence of peptides in the mass spectrometry data. PepQuery2 tool allows researchers to identify and validate known peptides in mass spectrometry-based proteomics datasets and a local protein sequence search database^37,38^. It generates outputs such as peptide-spectrum match (PSM) validation with confidence based on P-values and other features. For this confirmation, we subjected the 20,563 peptide sequences detected by any of the search algorithms to PepQuery analysis against our MS/MS data.

### Peptide assignment

Peptide reports from searches from all three algorithms were aligned to generate a master sheet. The peptides that were detected in the 2X background database search exclusively were subjected to BLAST-P analysis to ascertain that they were not present in the 30-organism database. Peptides that were detected in both the background and the 30-organism database were assigned to the 30-organism database.

### Quantitative Analysis

To generate quantitative data for analysis, the protein intensities from each organism were summed. The percentage of the total protein intensity was then compared to the expected percentage of each organism (Figure 5). Species abundances were calculated for each search condition by summing the unique protein intensities of individual species and dividing by the total protein intensity across all species within each replicate. These values were then averaged to determine the mean percentage abundances. Organisms were categorized based on their abundance levels: lowest (Ne1, F0, ES18, P22, Nm1, BXL, and Nu1), lower (BS, NV, 841, PaD, DVH, Am2, and 137), higher (KF7, CV, AK199, 259, HB2, VF, and PD), and highest (CRH, ATN, K12, Pfl, SMS, Cup, and LT2). The summed protein-level intensities for each abundance tier were compared to their expected percentages.

For peptide quantification, peptide intensities from proteins of each organism were summed. The abundance of each organism was represented as the sum intensity of all unique peptides annotated to that organism. The percentage of the total peptide intensity was then compared to the expected percentage for each organism. Species abundances were calculated for each search condition by summing the unique peptide intensities of individual species and dividing by the total peptide intensity across all species within each replicate. These values were then averaged to determine the mean percentage abundances. Organisms were categorized based on their abundance levels: lowest (Ne1, F0, ES18, P22, Nm1, BXL, and Nu1), lower (BS, NV, 841, PaD, DVH, Am2, and 137), higher (KF7, CV, AK199, 259, HB2, VF, and PD), and highest (CRH, ATN, K12, Pfl, SMS, Cup, and LT2). The summed peptide intensities for each abundance tier were compared to their expected percentages.

To mitigate issues related to protein inference, only peptides unambiguously assigned to a single organism were used to calculate species abundances within the microbiome community, following the methodology outlined in the original study for which these samples were generated^30^.

## RESULTS

To assess the performance of three software suites (MaxQuant, FragPipe, and Scribe) for quantitative metaproteomics, we analyzed the DDA-MS datasets generated from four biological replicates of the UNEVEN mock community (**Figure 1**)^30^. This UNEVEN mock community of 30 species was designed to cover a large range of species abundances, both at the level of cell number and proteinaceous biomass, to test for the dynamic range and detection limits of the quantification methods.

**Figure 1.**
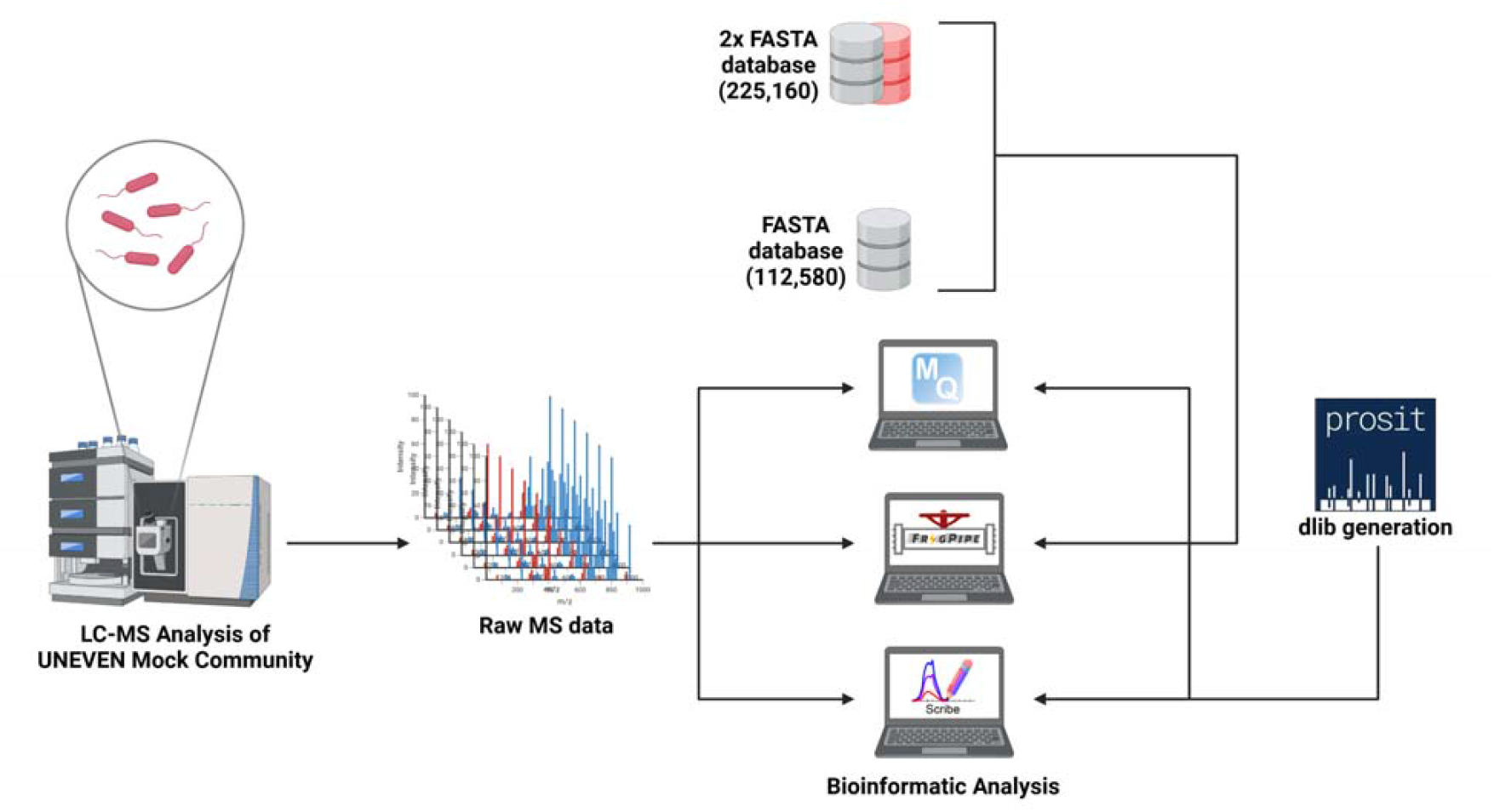
Experiment design overview. DDA-MS datasets were generated from four biological replicates of the UNEVEN mock community samples (Kleiner et al 2017), comprising microbial species that encompass a large dynamic range. The MS data was searched against the protein sequences database (1X database with 29 microorganisms; shown as a grey cylinder) along with a similar number of randomly selected protein sequences from the Integrated Gene Catalog Database (IGC, shown as a red cylinder; 2X database). The DDA-MS datasets were searched against the FASTA databases using two software suites (MaxQuant and FragPipe) for quantitative metaproteomics analyses. The DDA-MS datasets were searched against the spectral library and protein FASTA databases for the Scribe software.

Out of the 30 organisms (Supplementary Table 1), 28 organisms were detected in this study using all peptide intensities (**Figure 2**). Bacteriophages F2 and M13 were not detected in any of the searches. Low-abundance Phage P22 proteins were detected by all three search software algorithms (Supplementary Table 2). PepQuery analysis of the peptides associated with the phage P22 confirmed the presence of two peptides (NVLAQDATFSVVR and QVAGFDDVLR) in the mass spectrometry datasets. Low-abundance Phage F0 was detected by FragPipe and Scribe (**Figure 2**), and PepQuery analysis confirmed the presence of the F0 peptide (EVESITPDEIQG). Low-abundance Phage ES18 was only detected by Scribe. However, the peptide from Phage ES18 could not be confirmed using PepQuery (Supplementary Table 2). In all cases, the highest abundance organism LT2 was seen to have an observed percent abundance of approximately half its known percent abundance in the microbiome. Notably, the % CV values of Scribe were the lowest in cases where the organism was detected in all software suites (Figure 2B).

**Figure 2.**
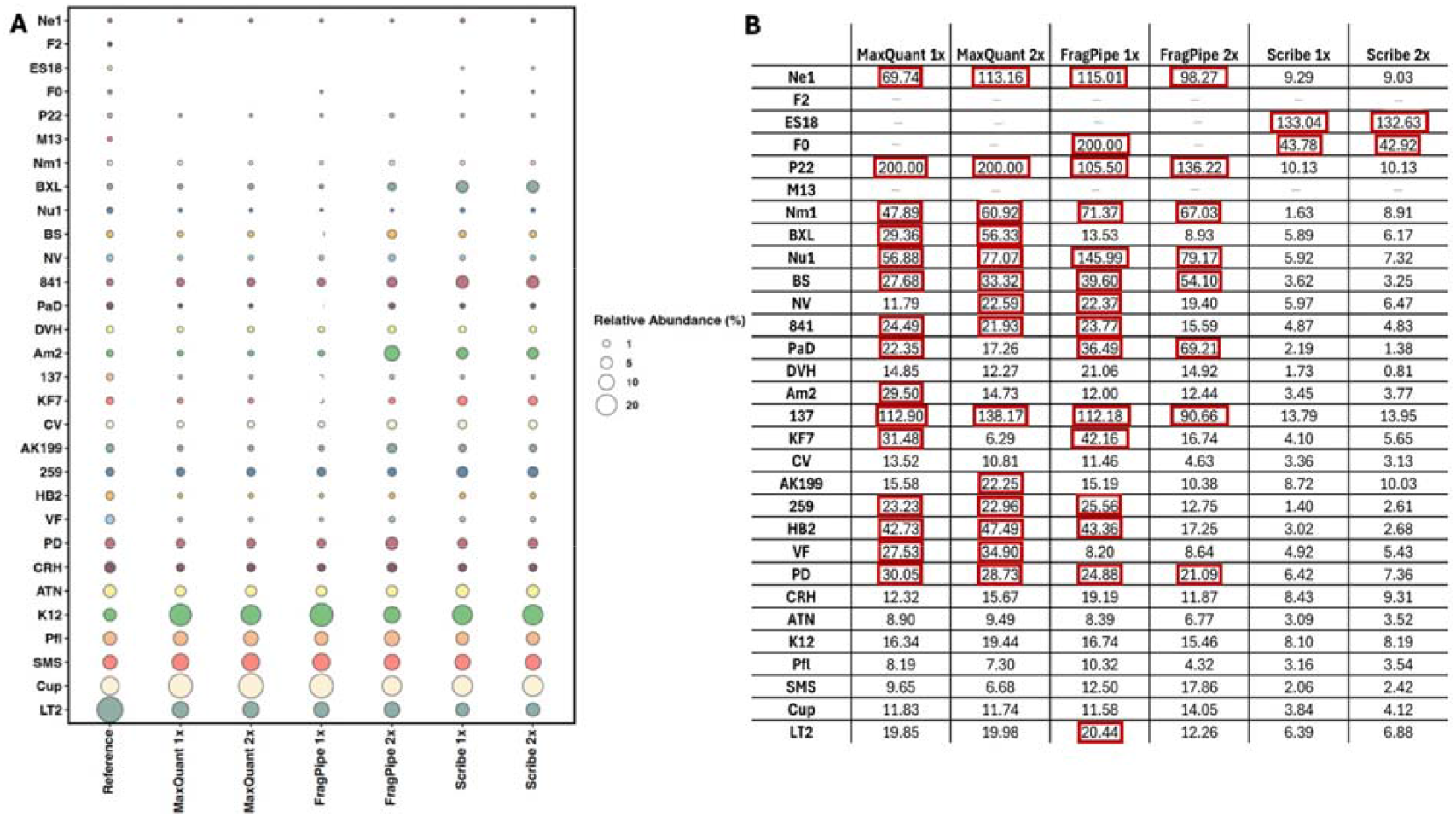
Abundances of organisms detected by MaxQuant, FragPipe, and Scribe. A) Diameter of the circles data points in the bubble plot represents the summed relative abundance of species-specific peptides for the species on the y-axis in each condition on the x-axis (variable analysis software with variable FASTA database size). The left-most column (Known % Abundance) shows the expected organismal abundance ranked from most abundant to the least. B) The matrix displays the percentage coefficient of variation (CV) for the DDA-MS measurements. CV values that are greater than 20% have been marked with red boxes.

Scribe detected the most number of proteins as compared to FragPipe and MaxQuant (**Figure 3A**). All the software identified a decreasing number of proteins and peptides with increasing database size (**Figure 3B**), and the number of background proteins and peptides (from the IGC database; **Figure 3**) increased as the database size increased from 1X to 2X. FragPipe and Scribe identified higher numbers of peptides and proteins as compared to MaxQuant. Scribe detected a higher number of proteins (18%) when compared to FragPipe, while FragPipe detected a higher number of peptides (2%) as compared to Scribe. The average number of peptides detected per protein were lower in Scribe versus FragPipe or MaxQuant (Supplementary Table 3). Moreover, Scribe had a substantially higher percentage of proteins detected with single peptides as compared to Fragpipe and MaxQuant.

**Figure 3.**
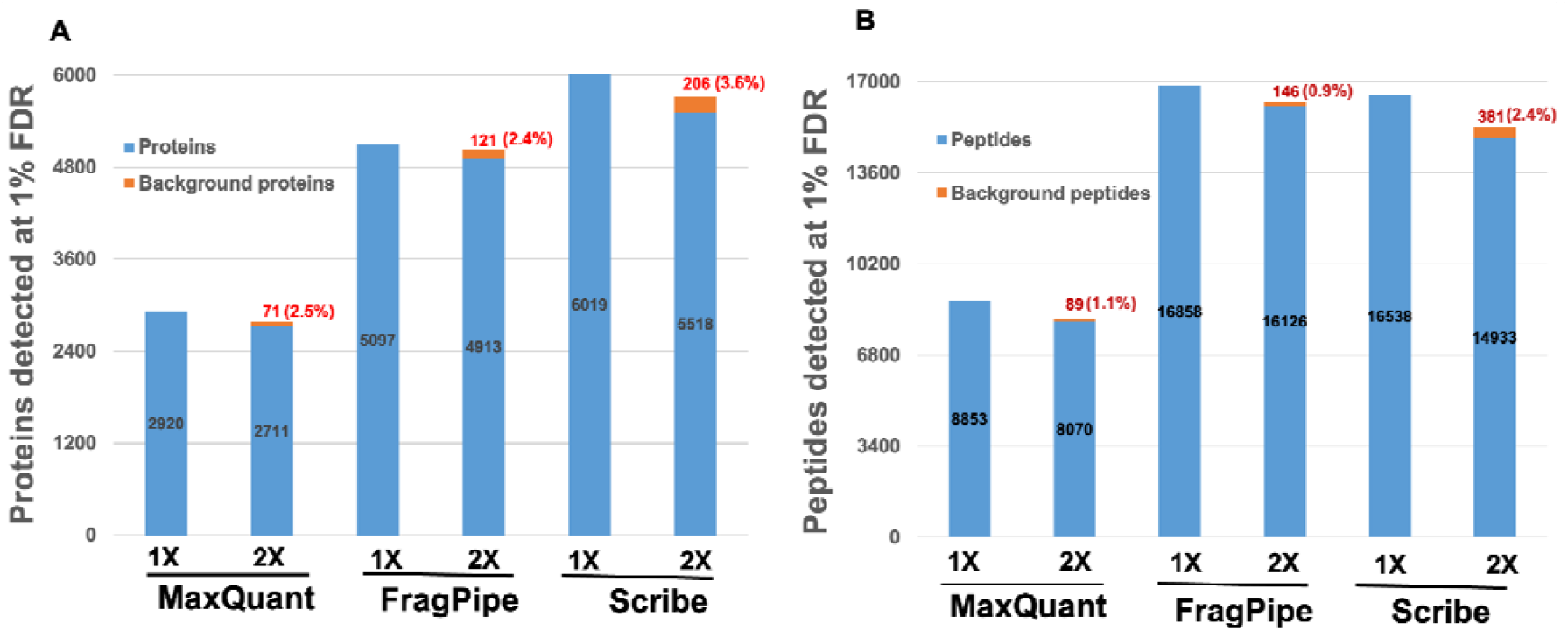
Proteins and peptides were detected from the synthetic microbiome database and the background (IGC database). **A)** Proteins detected at 1% FDR by MaxQuant, FragPipe, and Scribe. Detected proteins from the synthetic microbiome database are shown in blue, while background proteins are shown in red. 1X and 2X represent the increasing database size (See Methods and Figure 1). Background proteins were confirmed to ensure that the proteins did not include any proteins from the 30-organisms database. The percentage figures in parentheses indicate the percentage of background proteins detected in the 2X datasets. **B)** Peptides detected at 1% FDR by MaxQuant, FragPipe, and Scribe. Detected peptides from the synthetic microbiome database are in blue, while “background peptides” (peptide sequences of proteins likely not present in the sample) are shown in orange. Background peptides were confirmed by BLASTP analysis against proteins from the 30-organisms database. The percentage figures in parentheses indicate the percentage of background peptides detected in the 2X datasets.

FragPipe detected the most shared peptides across both datasets, as compared to Scribe and MaxQuant (**Supplementary Figure 1A**); Scribe was notable for having the highest percentage of unique peptides for each library when comparing the 1X and 2X database searches, suggesting that the use of a spectral library may allow for deeper, non-reproducible sequencing of the results. Moreover, the extent of overlap amongst the peptide detections from the three search algorithms can also be assessed by estimating the number of peptides detected exclusively by each of the search algorithms (**Supplementary Figure 1B**). FragPipe detected the highest number of unique peptides (for all database size searches) as compared to Scribe and then followed by MaxQuant. Unsurprisingly, increasing the database size with background proteins resulted in fewer shared peptides across the three platforms due to the potential for the stochastic “detection” of background proteins in the 2x library.

In terms of background proteins detected, FragPipe showed the best performance with the lowest percentage (2.4%) of background proteins detected (**Figure 3**, 2X database search). As compared to this, the other two software detected a higher percentage of background proteins, ranging from 2.5% (2X database search for MaxQuant) to 3.6% (2X database search for Scribe).

We investigated the nature of the peptides that match proteins from organisms that are not in the artificial community (termed as ‘background peptides’ in **Figure 3B and Figure 4**). We found that two background peptides (IINEPTAAALAYGLDKEVGNR, SGIAVGMATNIPPHNLR) were shared with proteins from the 30-organism database. After a BLASTP evaluation of these background peptides against the 30-microorganism database, we assigned these background peptides to the panel of 30-microorganism peptides if they matched with 100% identity. The background peptide detection rate ranged from 0.9% (for FragPipe; **see Figure 3B)** to 1.1% (for MaxQuant). Scribe had the highest background peptide detection rate of 2.5%.

**Figure 4.**
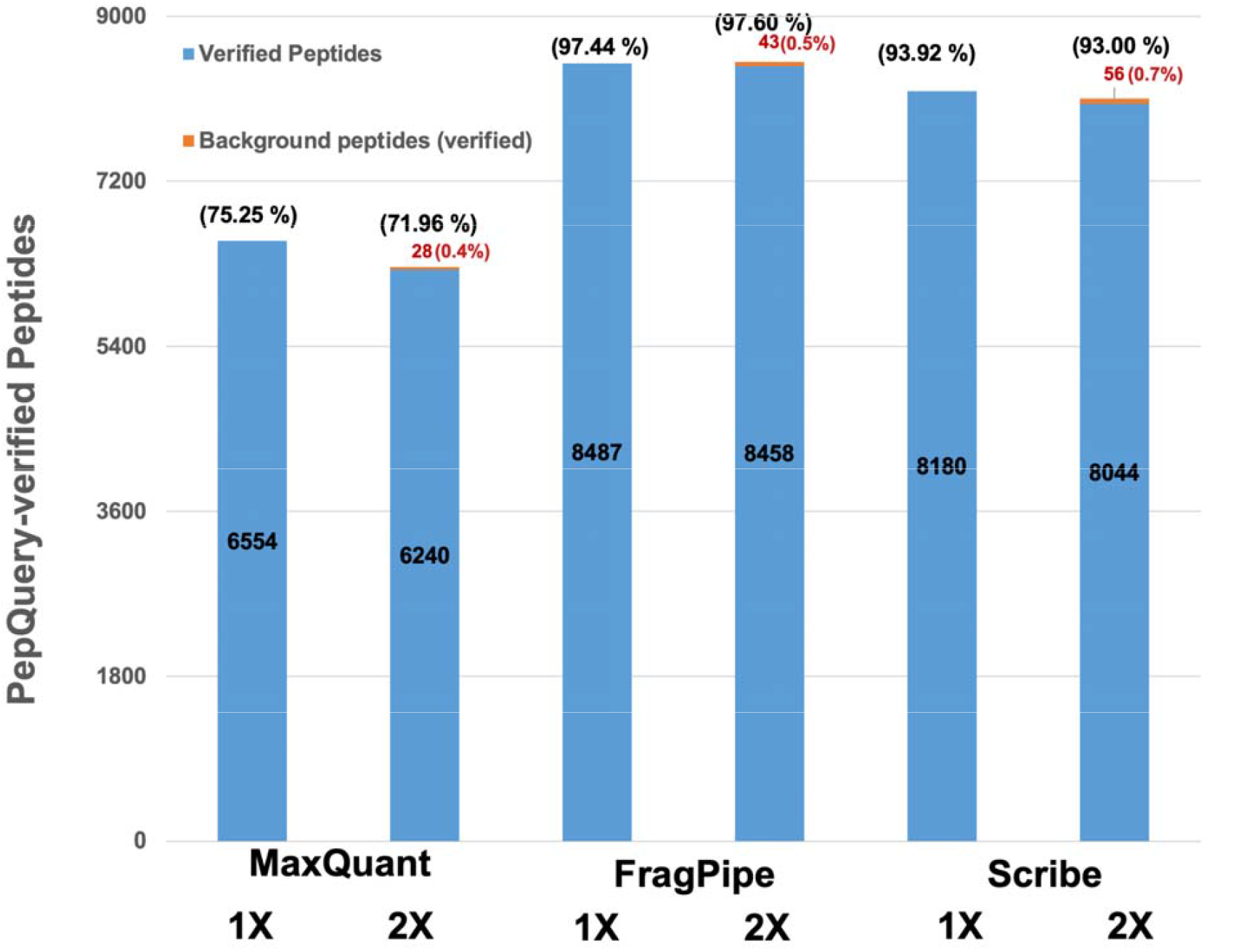
PepQuery analysis of detected peptides. 20,563 peptides that were detected by all three search algorithms against all search databases (1X and 2X) were subjected to PepQuery analysis. As a result, 8,710 peptides were confirmed to be present in the datasets. The figure shows the number of peptides detected by each search algorithm against variable database sizes. The numbers in blue-colored bars diagrams are assigned to synthetic microbiome proteins, while the numbers in red represent peptides that are assigned to background proteins. The percentages in parentheses (in black font) represent the percentage of 8,710 PepQuery-confirmed peptides detected in each search. The percentage figures in parentheses (in red font) indicate the percentage of background verified peptides detected in the 2X datasets.

Since most of the metaproteomics searches involve searches against large protein sequence databases, there is an increased chance of detection of false positives. Given that all of the modern MS-based proteomics studies use false discovery rate analysis, the confirmation of detected peptides by using orthogonal methods gains particular significance. A total of 8,710 peptides out of 20,563 peptides were verified to be present in the MS datasets using PepQuery, a peptide-centric database search engine.

For all three search algorithms, the number of verified peptides (based on PepQuery analysis) decreases as the database size increases (**Figure 4**). FragPipe showed the highest number of verified peptides (ranging from 8487 to 8501 peptides) with a verification percentage ranging from 97.4% to 97.6%. Scribe shows a verification percentage ranging from 93.9% to 93.0% (with verified peptides ranging from 8100 to 8180 peptides), and MaxQuant shows a lower verification percentage ranging from 71.9% to 75.2% (with verified peptides ranging from 6268 to 6554 peptides). The detection rates for background-verified peptides were 0.45% for MaxQuant, 0.52% for FragPipe, and 0.7% for Scribe. (**see Figure 4)**. After PepQuery analysis of all the peptides against the 30-microorganisms database, the background peptide detection rate for MaxQuant improved from 1.1% to 0.45% (**Figure 3B and Figure 4**), and the background peptide detection rate for FragPipe improved from 0.9% to 0.51%. Scribe, which had the highest background peptide detection rate of 2.49% showed an improvement to an impressive 0.69% after PepQuery analysis.

We assessed the quantitative accuracy of the three search algorithms by comparing the measured protein and peptide intensities to the expected abundance values based on the relative amount of the microorganisms making up the UNEVEN mock community and ascertaining the fold difference between these values as expressed via the 95% confidence intervals of these fold differences (**Figure 5**).

**Figure 5.**
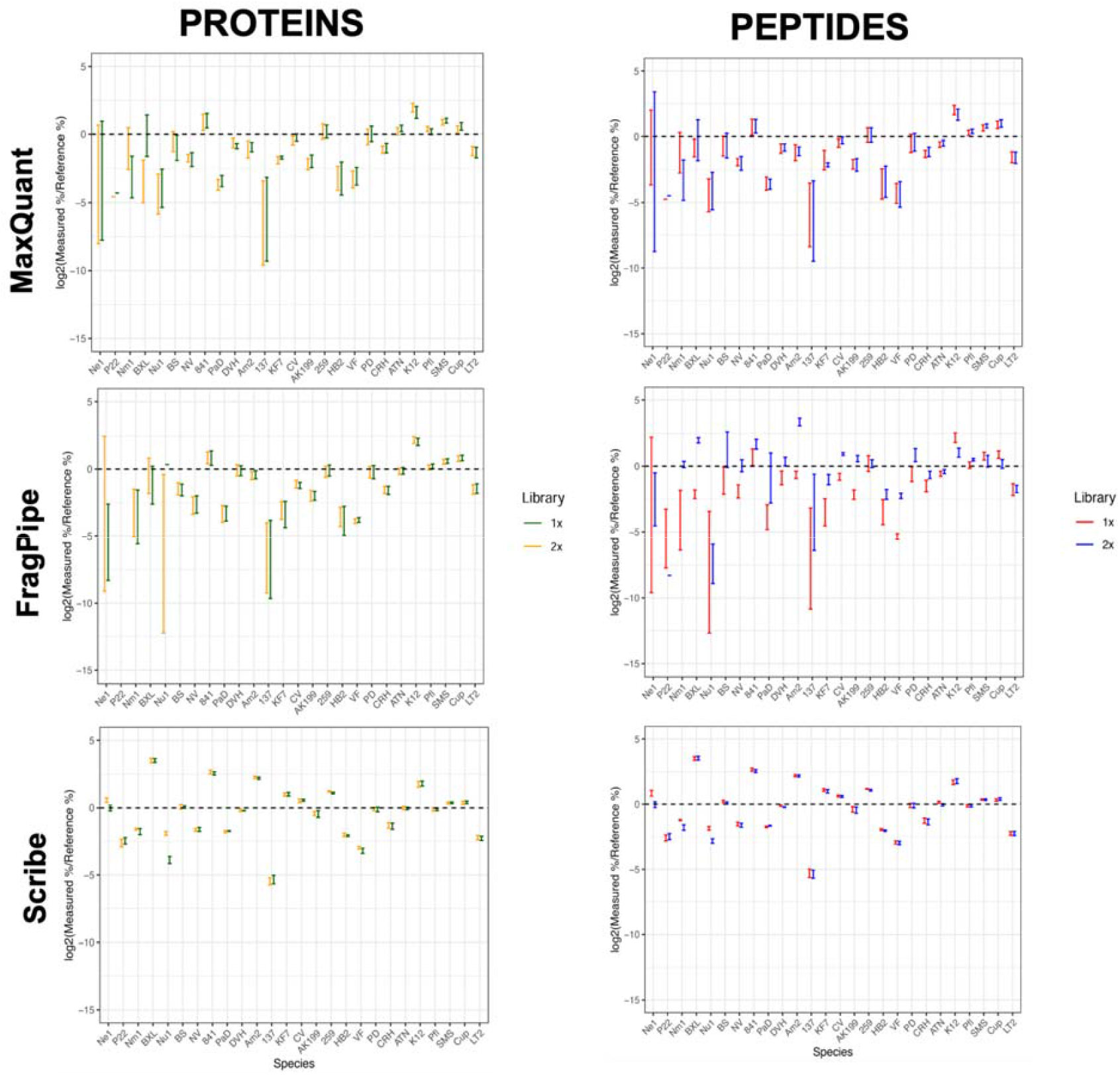
Quantitative analyses of the search results to demonstrate the relationship between measured and reference values in the Scribe platform. Box plots show the 95% confidence intervals of the fold difference between the measured and the reference % abundances of all species for the individual software suites. The line at 0.0 indicates the absence of any difference between expected and measured values.

For both MaxQuant and FragPipe, protein-level and peptide-level data showed very large confidence intervals for the lowest abundance organisms (**Figure 5**). These intervals become narrower for organisms of higher abundance and are minimal for the highest abundance organisms (**Supplementary Figure 2**). Additionally, the FragPipe peptide quantitative data reveal differences between the 1X and 2X searches, suggesting that the inclusion of background proteins in the database affects quantification when using the summed peptide intensities.

Quantitative analysis showed that Scribe had the most precise measurements of the species abundances, with the smallest confidence intervals of the species (**Figure 5**). For the lowest and lower abundance organisms, deviations from zero remained within 5-fold and improved with increasing organism abundance (**Supplementary Figure 2**). Scribe quantified higher abundance organisms with confidence intervals closest to zero, though all tools performed comparably for LT2, the highest abundance organism. Additionally, the data revealed minimal differences between the 1X and 2X searches for protein and peptide quantitation, indicating that Scribe’s quantification method is least affected by the inclusion of background proteins in the database.

## DISCUSSION

Most researchers who use MS-based metaproteomics in their work use DDA-MS methods for data generation. These DDA-MS-based metaproteomics workflows start with large database searches against public repository databases^39^ or metagenomics dataset-derived databases. These large database searches present challenges in low sensitivity detections and higher false-positive detections. Iterative database processing methods^18,20,21,34,40–42^ have been used in order to address this issue.

In this study, we evaluated the spectral library search algorithm Scribe along with other publicly available search algorithms such as MaxQuant and FragPipe for the identification, verification and quantification of microbial peptides and proteins from the organisms present in the UNEVEN mock community. We also estimated the detection of background peptides and proteins from the Integrated Gene Catalog (IGC) database. For this dataset, FragPipe performed best, followed by Scribe and MaxQuant, both in terms of detection of peptides and lower false positive rate at protein or peptide level. This is unsurprising, given the low false discovery proportions noted using FragPipe in a recent entrapment study by Wen *et al*.^43^

The background peptides that were detected by all of the search algorithms were analyzed by BLASTP analysis against the mock community database. The peptide-centric PepQuery tool was used to verify peptides detected from mass spectrometry-based datasets using all search algorithms. PepQuery is an alternative to traditional spectrum-centric search engines by focusing on specific peptides or proteins. Among the three algorithms, FragPipe showed the highest number of verified peptides detected with the highest verification percentage. It should be noted that MSFragger uses a scoring algorithm called Hyperscore to evaluate peptide-spectrum matches (PSMs) based on the quality of the match between experimental and theoretical spectra. PepQuery also uses Hyperscore to evaluate the quality of peptide-spectrum matches^44^. Additional investigation will be needed to determine if the increased number of PepQuery-verified peptides from FragPipe is due to the use of a similar scoring algorithm. Interestingly, the background peptide detection rate after the PepQuery verification step decreased, thus highlighting the utility of PepQuery as a tool for peptide verification. It was, however, surprising that some of these background peptides were verified by PepQuery, since we searched against the 30-organism database. Alternative PSM verification methods should characterize background peptides to assess whether detected peptides are genuine or the result of contamination. This characterization will help distinguish true peptide detections from false positives caused by incorrect matching of fragment spectra to peptide sequences.

Along with detection, we also evaluated the three search algorithms for the quantification of organisms added at known relative abundances. Scribe detected more low-abundance organisms (such as phages) from these samples. For quantification, MaxQuant uses the MaxLFQ algorithm^45^, FragPipe uses IonQuant^46^, and Scribe uses an interference rejection algorithm that draws inspiration from DIA-based quantification^27^. For quantitative accuracy, expressed as the confidence intervals of the log2 of the ratio of the detected versus known percentage abundances, Scribe performed best, followed by FragPipe and MaxQuant. Unlike FragPipe and MaxQuant, Scribe uses an interference rejection algorithm to quantify peptides, which we suspect may perform better in high-congestion samples with many co-eluting peptides. Simply described, this algorithm uses the calculated precursor isotopic distribution to constrain how much signal can be assigned to a peptide, which improves quantification accuracy when peptide intensities are low or missing. None of the search engines tested here were able to detect the phages M13 or F2; a previous analysis^47^ by some of the authors looking into the relative utility of DIA- and DDA-MS for metaproteomics was also unable to detect M13 proteins known to be present, suggesting that this phage may pose unique challenges in detection via mass spectrometry. Interestingly, in the previous Rajczewski *et al*. phage F2 was able to be detected in this microbiome by higher-end Orbitrap mass spectrometers (Fusion and Eclipse Tribrid Mass Spectrometers) than was used in this study, suggesting that increased resolution may be necessary for the detection of extremely low abundance organisms.

The three search platforms display usability trade-offs based on speed, sensitivity, accuracy, quantitation and large database search applications (**Supplementary Table 4**). MaxQuant, although widely used and stable, required longer runtimes and yielded fewer identifications and lowest proportion of PepQuery-verified peptide identifications, particularly with larger database search. FragPipe achieved the fastest runtime of the tested search engines and the highest proportion of PepQuery-verified peptide identifications even with larger database search. Other search engines, such as Sage^48^, have been shown to work even faster than FragPipe but were not tested in this study. Scribe showed sensitivity for low-abundance species and had the most reproducible quantitative measurements for all species but has limited usability for large database searches. Together, these findings should guide researchers in their open-source software selection for data-dependent MS searches based on the requirements for their metaproteomics searches.

One of the important aspects of metaproteomics studies is to characterize protein functions and taxonomy associated with the microbiome under study, based on protein identities output from the metaproteomic workflow. Since this information is based on the veracity of the underlying peptide detections, it was important to confirm the detection of microbial peptides. We used PepQuery for this analysis, although alternative methods such as spectral angles can also be used. For this dataset, we observed that FragPipe did the best with the detection of the PepQuery-confirmed peptides, followed by Scribe and MaxQuant.

We believe our evaluation offers metaproteomics researchers a valuable resource for selecting software for their DDA-MS searches based on the focus of their study. Whether prioritizing peptide detection, low error rates, sensitivity, accuracy, or confirmation of detected peptides, researchers can use our results, obtained with standard parameters, to choose the most suitable tool for their metaproteomics analysis.

## Acknowledgements/Funding

This work was funded by the National Institute of General Medical Sciences of the National Institutes of Health under Award Numbers R35GM138362 (MK) and R35GM150723 (BCS). The Proteogenomics Shared Resource in the University of Minnesota Masonic Cancer Center (NCI grant P30CA077598) provided additional support for S.M., P.D.J., and T.J.G. The European Research Council via ERC Starting Grant (#101077037) supported W.G. and M.W. We would like to thank Dechen Bhuming for her assistance in analysis of data for the earlier versions of the manuscript.

## Data Availability

Data can be accessed through Zenodo at https://zenodo.org/records/15392474. Raw mass spectrometry data are available at the PRIDE database under the entry PXD054415.

## Significance of the study

Metaproteomics requires a balance between high numbers of peptide and protein identification and confidence in the accuracy of the identifications made. We demonstrate the utility of the Scribe search engine for metaproteomics applications, as it was found to detect low-abundance proteins with accurate quantitation than other DDA-MS search engines. This tool has great utility for both novel metaproteomics studies as well as hypothesis-generating experiments using previously acquired open source proteomics raw data.

## References

(1) Salvato, F.; Hettich, R. L.; Kleiner, M. Five Key Aspects of Metaproteomics as a Tool to Understand Functional Interactions in Host-Associated Microbiomes. PLOS Pathog. 2021, 17 (2), e1009245. 10.1371/journal.ppat.1009245.

(2) Zhang, X.; Ning, Z.; Mayne, J.; Figeys, D. Clinical Microbiome Analysis by Mass Spectrometry-Based Metaproteomics. Annu. Rev. Anal. Chem. Palo Alto Calif 2025. 10.1146/annurev-anchem-071124-113819.

(3) Van Den Bossche, T.; Armengaud, J.; Benndorf, D.; Blakeley-Ruiz, J. A.; Brauer, M.; Cheng, K.; Creskey, M.; Figeys, D.; Grenga, L.; Griffin, T. J.; Henry, C.; Hettich, R. L.; Holstein, T.; Jagtap, P. D.; Jehmlich, N.; Kim, J.; Kleiner, M.; Kunath, B. J.; Malliet, X.; Martens, L.; Mehta, S.; Mesuere, B.; Ning, Z.; Tanca, A.; Uzzau, S.; Verschaffelt, P.; Wang, J.; Wilmes, P.; Zhang, X.; Zhang, X.; Li, L.; Initiative, T. M. The Microbiologist’s Guide to Metaproteomics. iMeta n/a (n/a), e70031. 10.1002/imt2.70031.

(4) Wilmes, P.; Heintz-Buschart, A.; Bond, P. L. A Decade of Metaproteomics: Where We Stand and What the Future Holds. Proteomics 2015, 15 (20), 3409–3417. 10.1002/pmic.201500183.

(5) Van Den Bossche, T.; Kunath, B. J.; Schallert, K.; Schäpe, S. S.; Abraham, P. E.; Armengaud, J.; Arntzen, M. Ø.; Bassignani, A.; Benndorf, D.; Fuchs, S.; Giannone, R. J.; Griffin, T. J.; Hagen, L. H.; Halder, R.; Henry, C.; Hettich, R. L.; Heyer, R.; Jagtap, P.; Jehmlich, N.; Jensen, M.; Juste, C.; Kleiner, M.; Langella, O.; Lehmann, T.; Leith, E.; May, P.; Mesuere, B.; Miotello, G.; Peters, S. L.; Pible, O.; Queiros, P. T.; Reichl, U.; Renard, B. Y.; Schiebenhoefer, H.; Sczyrba, A.; Tanca, A.; Trappe, K.; Trezzi, J.-P.; Uzzau, S.; Verschaffelt, P.; von Bergen, M.; Wilmes, P.; Wolf, M.; Martens, L.; Muth, T. Critical Assessment of MetaProteome Investigation (CAMPI): A Multi-Laboratory Comparison of Established Workflows. Nat. Commun. 2021, 12 (1), 7305. 10.1038/s41467-021-27542-8.

(6) Zhao, J.; Yang, Y.; Xu, H.; Zheng, J.; Shen, C.; Chen, T.; Wang, T.; Wang, B.; Yi, J.; Zhao, D.; Wu, E.; Qin, Q.; Xia, L.; Qiao, L. Data-Independent Acquisition Boosts Quantitative Metaproteomics for Deep Characterization of Gut Microbiota. Npj Biofilms Microbiomes 2023, 9 (1), 1–14. 10.1038/s41522-023-00373-9.

(7) Aakko, J.; Pietilä, S.; Suomi, T.; Mahmoudian, M.; Toivonen, R.; Kouvonen, P.; Rokka, A.; Hänninen, A.; Elo, L. L. Data-Independent Acquisition Mass Spectrometry in Metaproteomics of Gut Microbiota—Implementation and Computational Analysis. J. Proteome Res. 2020, 19 (1), 432–436. 10.1021/acs.jproteome.9b00606.

(8) Rajczewski, A. T.; Blakeley-Ruiz, J. A.; Meyer, A.; Vintila, S.; McIlvin, M. R.; Van Den Bossche, T.; Searle, B. C.; Griffin, T. J.; Saito, M. A.; Kleiner, M.; Jagtap, P. D. Data-Independent Acquisition Mass Spectrometry as a Tool for Metaproteomics: Interlaboratory Comparison Using a Model Microbiome. Proteomics 2025, e202400187. 10.1002/pmic.202400187.

(9) Eng, J. K.; Searle, B. C.; Clauser, K. R.; Tabb, D. L. A Face in the Crowd: Recognizing Peptides Through Database Search*. Mol. Cell. Proteomics 2011, 10 (11), R111.009522. 10.1074/mcp.R111.009522.

(10) Marcotte, E. M. How Do Shotgun Proteomics Algorithms Identify Proteins? Nat. Biotechnol. 2007, 25 (7), 755–757. 10.1038/nbt0707-755.

(11) Blakeley-Ruiz, J. A.; Kleiner, M. Considerations for Constructing a Protein Sequence Database for Metaproteomics. Comput. Struct. Biotechnol. J. 2022, 20, 937–952. 10.1016/j.csbj.2022.01.018.

(12) Griss, J. Spectral Library Searching in Proteomics. PROTEOMICS 2016, 16 (5), 729–740. 10.1002/pmic.201500296.

(13) Nowatzky, Y.; Benner, P.; Reinert, K.; Muth, T. Mistle: Bringing Spectral Library Predictions to Metaproteomics with an Efficient Search Index. Bioinformatics 2023, 39 (6), btad376. 10.1093/bioinformatics/btad376.

(14) Yang, Y.; Liu, X.; Shen, C.; Lin, Y.; Yang, P.; Qiao, L. In Silico Spectral Libraries by Deep Learning Facilitate Data-Independent Acquisition Proteomics. Nat. Commun. 2020, 11 (1), 146. 10.1038/s41467-019-13866-z.

(15) Zhang, X.; Li, Y.; Shao, W.; Lam, H. Understanding the Improved Sensitivity of Spectral Library Searching over Sequence Database Searching in Proteomics Data Analysis. PROTEOMICS 2011, 11 (6), 1075–1085. 10.1002/pmic.201000492.

(16) VerBerkmoes, N. C.; Denef, V. J.; Hettich, R. L.; Banfield, J. F. Functional Analysis of Natural Microbial Consortia Using Community Proteomics. Nat. Rev. Microbiol. 2009, 7 (3), 196–205. 10.1038/nrmicro2080.

(17) Muth, T.; Benndorf, D.; Reichl, U.; Rapp, E.; Martens, L. Searching for a Needle in a Stack of Needles: Challenges in Metaproteomics Data Analysis. Mol. Biosyst. 2013, 9 (4), 578–585. 10.1039/C2MB25415H.

(18) Bassignani, A.; Plancade, S.; Berland, M.; Blein-Nicolas, M.; Guillot, A.; Chevret, D.; Moritz, C.; Huet, S.; Rizkalla, S.; Clément, K.; Doré, J.; Langella, O.; Juste, C. Benefits of Iterative Searches of Large Databases to Interpret Large Human Gut Metaproteomic Data Sets. J. Proteome Res. 2021, 20 (3), 1522–1534. 10.1021/acs.jproteome.0c00669.

(19) Zhang, X.; Ning, Z.; Mayne, J.; Moore, J. I.; Li, J.; Butcher, J.; Deeke, S. A.; Chen, R.; Chiang, C.-K.; Wen, M.; Mack, D.; Stintzi, A.; Figeys, D. MetaPro-IQ: A Universal Metaproteomic Approach to Studying Human and Mouse Gut Microbiota. Microbiome 2016, 4 (1), 31. 10.1186/s40168-016-0176-z.

(20) Kumar, P.; Johnson, J. E.; Easterly, C.; Mehta, S.; Sajulga, R.; Nunn, B.; Jagtap, P. D.; Griffin, T. J. A Sectioning and Database Enrichment Approach for Improved Peptide Spectrum Matching in Large, Genome-Guided Protein Sequence Databases. J. Proteome Res. 2020, 19 (7), 2772–2785. 10.1021/acs.jproteome.0c00260.

(21) Jagtap, P.; Goslinga, J.; Kooren, J. A.; McGowan, T.; Wroblewski, M. S.; Seymour, S. L.; Griffin, T. J. A Two-Step Database Search Method Improves Sensitivity in Peptide Sequence Matches for Metaproteomics and Proteogenomics Studies. PROTEOMICS 2013, 13 (8), 1352–1357. 10.1002/pmic.201200352.

(22) Potgieter, M. G.; Nel, A. J. M.; Fortuin, S.; Garnett, S.; Wendoh, J. M.; Tabb, D. L.; Mulder, N. J.; Blackburn, J. M. MetaNovo: An Open-Source Pipeline for Probabilistic Peptide Discovery in Complex Metaproteomic Datasets. PLOS Comput. Biol. 2023, 19 (6), e1011163. 10.1371/journal.pcbi.1011163.

(23) Muth, T.; Renard,Bernhard Y.; and Martens, L. Metaproteomic Data Analysis at a Glance: Advances in Computational Microbial Community Proteomics. Expert Rev. Proteomics 2016, 13 (8), 757–769. 10.1080/14789450.2016.1209418.

(24) Rechenberger, J.; Samaras, P.; Jarzab, A.; Behr, J.; Frejno, M.; Djukovic, A.; Sanz, J.; González-Barberá, E. M.; Salavert, M.; López-Hontangas, J. L.; Xavier, K. B.; Debrauwer, L.; Rolain, J.-M.; Sanz, M.; Garcia-Garcera, M.; Wilhelm, M.; Ubeda, C.; Kuster, B. Challenges in Clinical Metaproteomics Highlighted by the Analysis of Acute Leukemia Patients with Gut Colonization by Multidrug-Resistant Enterobacteriaceae. Proteomes 2019, 7 (1), 2. 10.3390/proteomes7010002.

(25) Deutsch, E. W.; Perez-Riverol, Y.; Chalkley, R. J.; Wilhelm, M.; Tate, S.; Sachsenberg, T.; Walzer, M.; Käll, L.; Delanghe, B.; Böcker, S.; Schymanski, E. L.; Wilmes, P.; Dorfer, V.; Kuster, B.; Volders, P.-J.; Jehmlich, N.; Vissers, J. P. C.; Wolan, D. W.; Wang, A. Y.; Mendoza, L.; Shofstahl, J.; Dowsey, A. W.; Griss, J.; Salek, R. M.; Neumann, S.; Binz, P.-A.; Lam, H.; Vizcaíno, J. A.; Bandeira, N.; Röst, H. Expanding the Use of Spectral Libraries in Proteomics. J. Proteome Res. 2018, 17 (12), 4051–4060. 10.1021/acs.jproteome.8b00485.

(26) Gessulat, S.; Schmidt, T.; Zolg, D. P.; Samaras, P.; Schnatbaum, K.; Zerweck, J.; Knaute, T.; Rechenberger, J.; Delanghe, B.; Huhmer, A.; Reimer, U.; Ehrlich, H.-C.; Aiche, S.; Kuster, B.; Wilhelm, M. Prosit: Proteome-Wide Prediction of Peptide Tandem Mass Spectra by Deep Learning. Nat. Methods 2019, 16 (6), 509–518. 10.1038/s41592-019-0426-7.

(27) Searle, B. C.; Shannon, A. E.; Wilburn, D. B. Scribe: Next Generation Library Searching for DDA Experiments. J. Proteome Res. 2023, 22 (2), 482–490. 10.1021/acs.jproteome.2c00672.

(28) Cox, J.; Mann, M. MaxQuant Enables High Peptide Identification Rates, Individualized p.p.b.-Range Mass Accuracies and Proteome-Wide Protein Quantification. Nat. Biotechnol. 2008, 26 (12), 1367–1372. 10.1038/nbt.1511.

(29) Kong, A. T.; Leprevost, F. V.; Avtonomov, D. M.; Mellacheruvu, D.; Nesvizhskii, A. I. MSFragger: Ultrafast and Comprehensive Peptide Identification in Shotgun Proteomics. Nat. Methods 2017, 14 (5), 513–520. 10.1038/nmeth.4256.

(30) Kleiner, M.; Thorson, E.; Sharp, C. E.; Dong, X.; Liu, D.; Li, C.; Strous, M. Assessing Species Biomass Contributions in Microbial Communities via Metaproteomics. Nat. Commun. 2017, 8 (1), 1558. 10.1038/s41467-017-01544-x.

(31) Wiśniewski, J. R.; Zougman, A.; Nagaraj, N.; Mann, M. Universal Sample Preparation Method for Proteome Analysis. Nat. Methods 2009, 6 (5), 359–362. 10.1038/nmeth.1322.

(32) Li, W.; Godzik, A. Cd-Hit: A Fast Program for Clustering and Comparing Large Sets of Protein or Nucleotide Sequences. Bioinforma. Oxf. Engl. 2006, 22 (13), 1658–1659. 10.1093/bioinformatics/btl158.

(33) Li, J.; Jia, H.; Cai, X.; Zhong, H.; Feng, Q.; Sunagawa, S.; Arumugam, M.; Kultima, J. R.; Prifti, E.; Nielsen, T.; Juncker, A. S.; Manichanh, C.; Chen, B.; Zhang, W.; Levenez, F.; Wang, J.; Xu, X.; Xiao, L.; Liang, S.; Zhang, D.; Zhang, Z.; Chen, W.; Zhao, H.; Al-Aama, J. Y.; Edris, S.; Yang, H.; Wang, J.; Hansen, T.; Nielsen, H. B.; Brunak, S.; Kristiansen, K.; Guarner, F.; Pedersen, O.; Doré, J.; Ehrlich, S. D.; MetaHIT Consortium; Bork, P.; Wang, J.; MetaHIT Consortium. An Integrated Catalog of Reference Genes in the Human Gut Microbiome. Nat. Biotechnol. 2014, 32 (8), 834–841. 10.1038/nbt.2942.

(34) Picciani, M.; Gabriel, W.; Giurcoiu, V.-G.; Shouman, O.; Hamood, F.; Lautenbacher, L.; Jensen, C. B.; Müller, J.; Kalhor, M.; Soleymaniniya, A.; Kuster, B.; The, M.; Wilhelm, M. Oktoberfest: Open-Source Spectral Library Generation and Rescoring Pipeline Based on Prosit. Proteomics 2024, 24 (8), e2300112. 10.1002/pmic.202300112.

(35) Wilhelm, M.; Zolg, D. P.; Graber, M.; Gessulat, S.; Schmidt, T.; Schnatbaum, K.; Schwencke-Westphal, C.; Seifert, P.; de Andrade Krätzig, N.; Zerweck, J.; Knaute, T.; Bräunlein, E.; Samaras, P.; Lautenbacher, L.; Klaeger, S.; Wenschuh, H.; Rad, R.; Delanghe, B.; Huhmer, A.; Carr, S. A.; Clauser, K. R.; Krackhardt, A. M.; Reimer, U.; Kuster, B. Deep Learning Boosts Sensitivity of Mass Spectrometry-Based Immunopeptidomics. Nat. Commun. 2021, 12, 3346. 10.1038/s41467-021-23713-9.

(36) Hulsen, T. DeepVenn -- a Web Application for the Creation of Area-Proportional Venn Diagrams Using the Deep Learning Framework Tensorflow.Js. arXiv September 27, 2022. 10.48550/arXiv.2210.04597.

(37) Wen, B.; Wang, X.; Zhang, B. PepQuery Enables Fast, Accurate, and Convenient Proteomic Validation of Novel Genomic Alterations. Genome Res. 2019, 29 (3), 485–493. 10.1101/gr.235028.118.

(38) Wen, B.; Zhang, B. PepQuery2 Democratizes Public MS Proteomics Data for Rapid Peptide Searching. Nat. Commun. 2023, 14, 2213. 10.1038/s41467-023-37462-4.

(39) Muth, T.; Kolmeder, C. A.; Salojärvi, J.; Keskitalo, S.; Varjosalo, M.; Verdam, F. J.; Rensen, S. S.; Reichl, U.; de Vos, W. M.; Rapp, E.; Martens, L. Navigating through Metaproteomics Data: A Logbook of Database Searching. Proteomics 2015, 15 (20), 3439–3453. 10.1002/pmic.201400560.

(40) Kertesz-Farkas, A.; Keich, U.; Noble, W. S. Tandem Mass Spectrum Identification via Cascaded Search. J. Proteome Res. 2015, 14 (8), 3027–3038. 10.1021/pr501173s.

(41) Stamboulian, M.; Li, S.; Ye, Y. Using High-Abundance Proteins as Guides for Fast and Effective Peptide/Protein Identification from Human Gut Metaproteomic Data. Microbiome 2021, 9 (1), 80. 10.1186/s40168-021-01035-8.

(42) Rooijers, K.; Kolmeder, C.; Juste, C.; Doré, J.; de Been, M.; Boeren, S.; Galan, P.; Beauvallet, C.; de Vos, W. M.; Schaap, P. J. An Iterative Workflow for Mining the Human Intestinal Metaproteome. BMC Genomics 2011, 12, 6. 10.1186/1471-2164-12-6.

(43) Wen, B.; Freestone, J.; Riffle, M.; MacCoss, M. J.; Noble, W. S.; Keich, U. Assessment of False Discovery Rate Control in Tandem Mass Spectrometry Analysis Using Entrapment. Nat. Methods 2025, 22 (7), 1454–1463. 10.1038/s41592-025-02719-x.

(44) Wen, B.; Wang, X.; Zhang, B. PepQuery Enables Fast, Accurate, and Convenient Proteomic Validation of Novel Genomic Alterations. Genome Res. 2019, 29 (3), 485–493. 10.1101/gr.235028.118.

(45) Cox, J.; Hein, M. Y.; Luber, C. A.; Paron, I.; Nagaraj, N.; Mann, M. Accurate Proteome-Wide Label-Free Quantification by Delayed Normalization and Maximal Peptide Ratio Extraction, Termed MaxLFQ. Mol. Cell. Proteomics MCP 2014, 13 (9), 2513–2526. 10.1074/mcp.M113.031591.

(46) Yu, F.; Haynes, S. E.; Nesvizhskii, A. I. IonQuant Enables Accurate and Sensitive Label-Free Quantification With FDR-Controlled Match-Between-Runs. Mol. Cell. Proteomics MCP 2021, 20, 100077. 10.1016/j.mcpro.2021.100077.

(47) Rajczewski, A. T.; Blakeley-Ruiz, J. A.; Meyer, A.; Vintila, S.; McIlvin, M. R.; Van Den Bossche, T.; Searle, B. C.; Griffin, T. J.; Saito, M. A.; Kleiner, M.; Jagtap, P. D. Data-Independent Acquisition Mass Spectrometry as a Tool for Metaproteomics: Interlaboratory Comparison Using a Model Microbiome. PROTEOMICS 2025, 25 (9–10), e202400187. 10.1002/pmic.202400187.

(48) Lazear, M. R. Sage: An Open-Source Tool for Fast Proteomics Searching and Quantification at Scale. J. Proteome Res. 2023, 22 (11), 3652–3659. 10.1021/acs.jproteome.3c00486.

