## Supplemental Information for "Comparative Performance of Scribe and Database Search Engines in Metaproteomic Profiling of a Ground-Truth Microbiome Dataset"

**Supplement S1 Table S1:** Protein sequence database components used in the proteomic analyses of composite microbiome samples

| **Label** | **Species** | **Protein Sequences Accession Number(s)** | **Input Protein Amount (ug)** | **Percentage** |
| --- | --- | --- | --- | --- |
| Ne1 | *Nitrosomonas europaeae* | UP000001416 | 60.58 | 0.082 |
| F2 | Phage F2 | UP000002127 | 62.04 | 0.084 |
| F0 | Phage F0 | UP000009070 | 65.28 | 0.088 |
| ES18 | Phage ES18 | UP000000970 | 65.55 | 0.088 |
| P22 | Phage P22 | UP000007960 | 78.77 | 0.106 |
| M13 | Phage M13 | UP000002111 | 109.02 | 0.147 |
| Nm1 | *Nitrosospira multiformis* | UP000002718 | 155.16 | 0.209 |
| BXL | *Burkholderia xenovorans* | UP000001817 | 321.37 | 0.433 |
| Nu1 | *Nitrosomonas ureae* | UP000056699 | 402.68 | 0.543 |
| BS | *Bacillus subtilis* | UP000001570 | 583.83 | 0.788 |
| NV | *Nitrososphaera viennensis* | GCA_000698785.1 | 607.41 | 0.819 |
| 841 | *Rhizobium leguminosarum bv. viciae* 3841 | UP000006575 | 680.47 | 0.918 |
| PaD | *Paracoccus denitrificans* | Used RAST to generate protein sequences. Available in the protein sequence database on PRIDE PXD006118 | 683.66 | 0.922 |
| DVH | *Desulfovibrio vulgaris* | UP000002194 | 701.58 | 0.946 |
| Am2 | *Alteromonas macleodii* | UP000006296 | 707.47 | 0.954 |
| 137 | *Staphylococcus aureus* ATCC 13709 | Used RAST to generate protein sequences. Available in the protein sequence database on PRIDE PXD006118 | 715.15 | 0.965 |
| KF7 | *Pseudomonas pseudoalcaligenes* | GCA_000262065.3 | 863.80 | 1.165 |
| CV | *Chromobacterium violaceum* | Used RAST to generate protein sequences. Available in the protein sequence database on PRIDE PXD006118 | 933.08 | 1.259 |
| AK199 | *Roseobacter sp.* AK199 | Used RAST to generate protein sequences. Available in the protein sequence database on PRIDE PXD006118 | 1183.43 | 1.596 |
| 259 | *Staphylococcus aureus* ATCC 25923 | GCA_000756205.1 | 1216.31 | 1.641 |
| HB2 | *Thermus Thermophilus* | UP000000592 | 1245.63 | 1.680 |
| VF | *Rhizobium leguminosarum bv. viciae* VF39 | GCA_000427765.1 | 1671.03 | 2.254 |
| PD | *Pseudomonas denitrificans* | UP000012082 | 2128.51 | 2.871 |
| CRH | *Chlamydomonas reinhardtii* | GCF_000002595.1 | 2962.73 | 3.996 |
| ATN | *Agrobacterium tumefaciens* | UP00000813 | 4186.28 | 5.647 |
| K12 | *Escherichia coli* | UP000000625 | 4290.54 | 5.788 |
| Pfl | *Pseudomonas fluorescens* | 2617270901 (IMG) | 4964.22 | 6.696 |
| SMS | *Stenotrophomonas maltophilia* | GCF_000613205.1 | 5946.27 | 8.021 |
| Cup | *Cupriavidus metallidurans* | UP000002429 | 11504.98 | 15.519 |
| LT2 | *Salmonella enterica typhimurium* (3 strains combined) | UP000001014 | 25037.78 | 33.773 |

**Supplement S1 Table S2:** Detection of phage peptides and their verification. Bacteriophage names are mentioned under the “Organism” column; Peptide sequences under the “Peptide” column; 1X and 2X refer to database sizes under the search algorithm columns. U1 to U4 under the “PepQuery” column refer to “Uneven” replicates. Yes refers to either detection (for search algorithms) and verification (under PepQuery).

|  |  | **MaxQuant** | | **FragPipe** | | **Scribe** | | **PepQuery** | | | |
| --- | --- | --- | --- | --- | --- | --- | --- | --- | --- | --- | --- |
| **Organism** | **Peptide** | **1X** | **2X** | **1X** | **2X** | **1X** | **2X** | **U1** | **U2** | **U3** | **U4** |
| **F0** | EVESITPDEIQGVR |  |  | **Yes** | **Yes** | **Yes** | **Yes** |  |  | **Yes** |  |
| **ES18** | DQEVGMEIVNDLIGVQTVLPVGK |  |  |  |  | **Yes** | **Yes** |  |  |  |  |
| **P22** | NVLAQDATFSVVR | **Yes** | **Yes** | **Yes** | **Yes** | **Yes** | **Yes** | **Yes** | **Yes** | **Yes** |  |
| **P22** | YTPPAASMQR |  |  |  |  | **Yes** | **Yes** |  |  |  |  |
| **P22** | ADDLRDETAYR |  |  | **Yes** | **Yes** |  |  |  |  |  |  |
| **P22** | QVAGFDDVLR |  |  | **Yes** |  |  |  |  |  |  | **Yes** |
| **P22** | ALNEGQIVTLAVDEIIETISAITPMAQK |  |  | **Yes** | **Yes** |  |  |  |  |  |  |
| **P22** | TTSFSIPDVGLNGIFATQGDISTLSGLCR |  |  | **Yes** | **Yes** |  |  |  |  |  |  |
| **P22** | VVDGTHVEITPKPVALDDVSLSPEQR |  |  | **Yes** | **Yes** |  |  |  |  |  |  |

**Supplement S1 Table S3:** Detection of proteins with single peptides and their percentages. Proteins detected with a single peptide were enlisted for all searches. Average number of peptides per protein were calculated.

| **Search** | **Proteins detected** | **Proteins detected with a single peptide** | **Percentage of single-peptide protein detections** | **Average number of peptides/protein** |
| --- | --- | --- | --- | --- |
| **MQ_1X** | 2957 | 1114 | 37.7 | 3.29 |
| **MQ_2X** | 2818 | 1090 | 38.7 | 3.19 |
| **FP_1X** | 5101 | 1820 | 35.7 | 3.63 |
| **FP_2X** | 5038 | 1850 | 36.7 | 3.61 |
| **Scribe_1X** | 6024 | 2826 | 46.9 | 2.94 |
| **Scribe_2X** | 5729 | 2724 | 47.5 | 2.85 |

**Supplement S1 Table S4:** Comparison of metaproteomics search software performance. Search times were measured using the same computational setup. Usability features such as speed, sensitivity, accuracy, quantitation and large database search applications are summarized.

| **Software** | **Search Time (Speed)** | **Detection sensitivity** | **Peptide detection veracity** | **Quantitative accuracy** | **Large database search application** |
| --- | --- | --- | --- | --- | --- |
| **MaxQuant** | 1 hr 2 mins | Lowest number of peptides and proteins detected. | Lowest (~75%) based on PepQuery verification. | Low quantitative accuracy for lowest, lower and higher abundance organisms. | Low detection, low detection accuracy, low uantitative accuracy in large database search. |
| **FragPipe** | 11 min | Highest number of peptides detected and intermediate number of proteins detected. | Highest (~97%) based on PepQuery verification. | Low quantitative accuracy for lowest, lower and higher abundance organisms. | Highest detection, highest detection accuracy, low uantitative accuracy in large database search. |
| **Scribe** | 1 hr 37 mins* | Intermediate number of peptides detected and highest number of proteins detected. | Intermediate (~93%) based on PepQuery verification. | High quantitative accuracy for lowest, lower and higher abundance organisms. | High detection, high detection accuracy, highest quantitaive accuracy in large database search. |

**On a resource with comparable RAM and higher number of cores.*

**A**

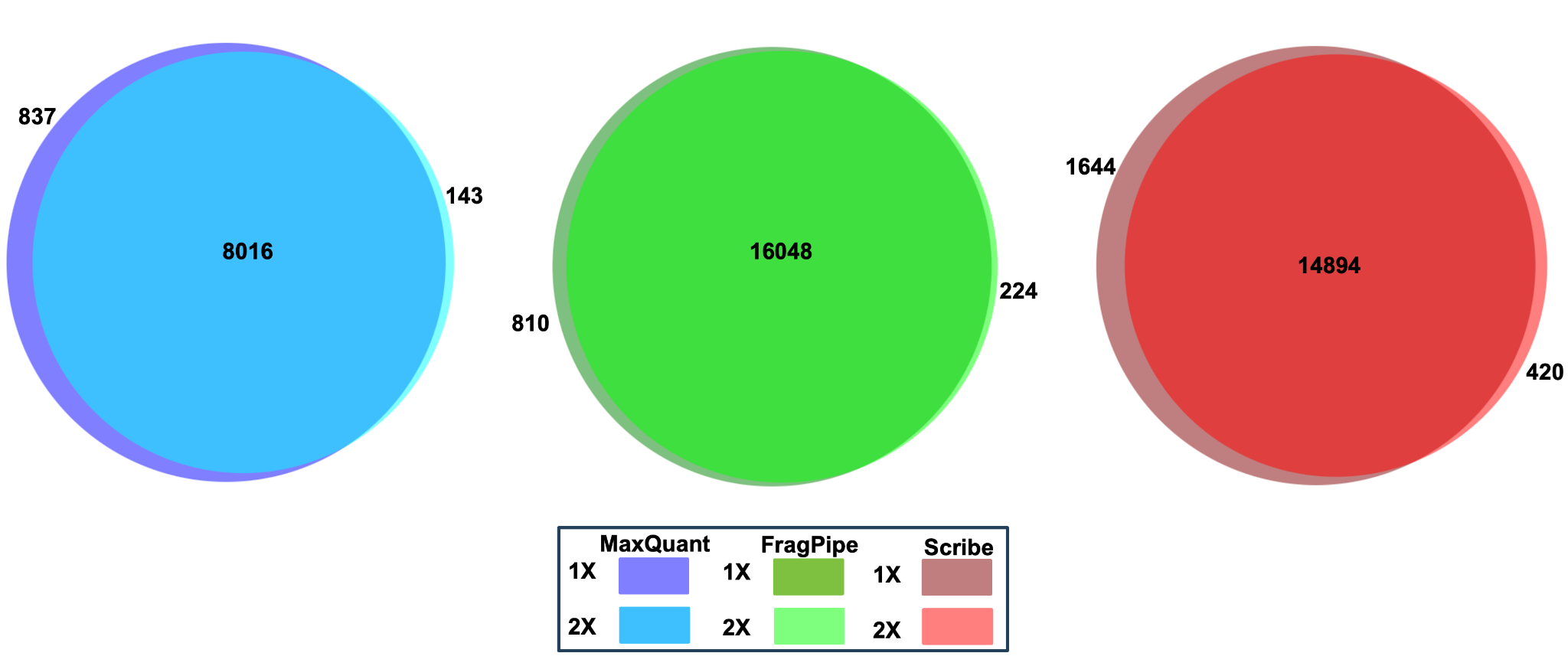

**B**

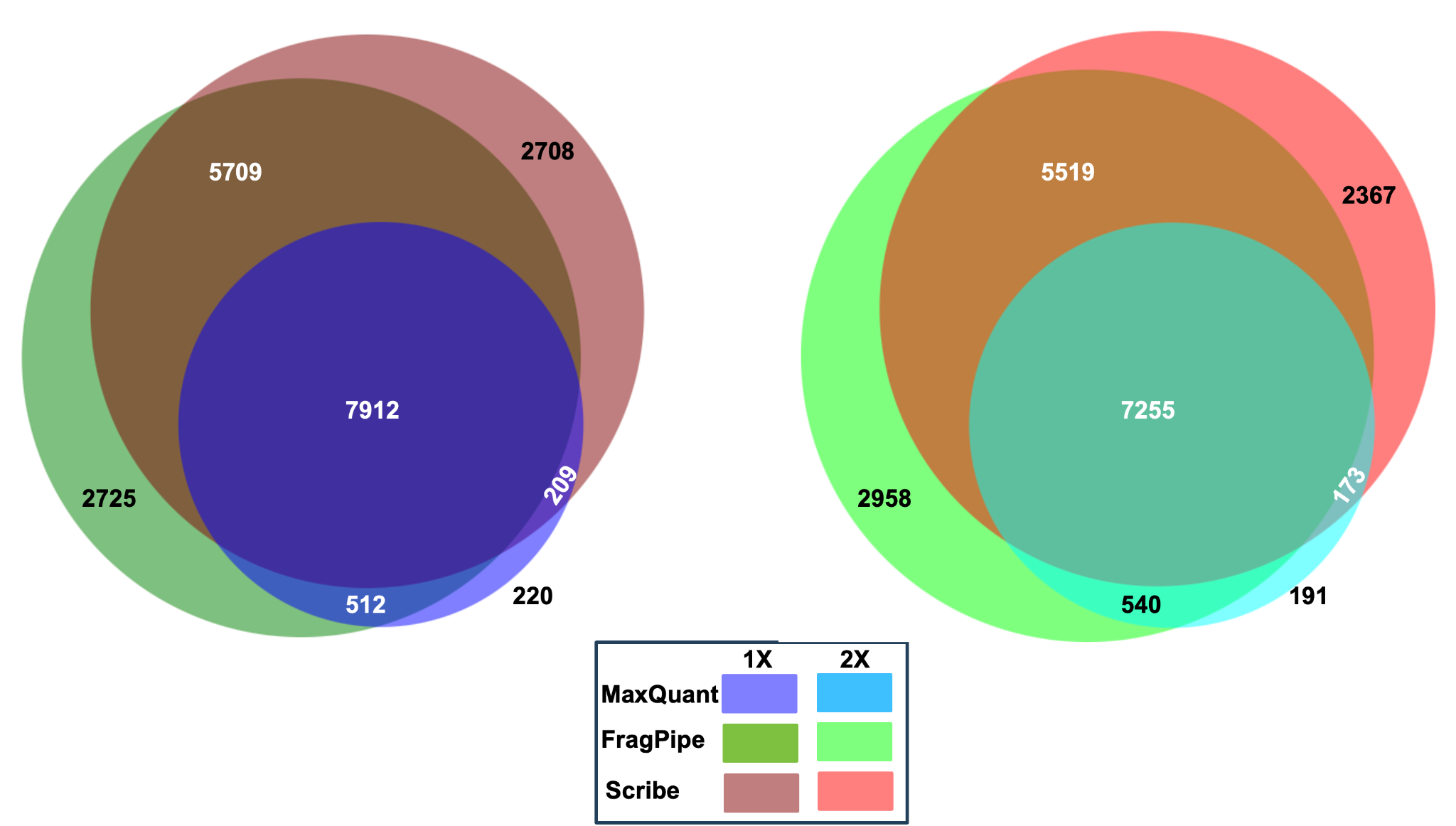
**Supplement S1 Figure S1. Overlap of peptides detected from the synthetic microbiome database. A)** Peptides detected by MaxQuant, FragPipe, and Scribe have been represented. The color scheme for the searches against databases of variable sizes. **B)** Peptides detected by 1X and 2X database size by the three search algorithms have been represented. The color scheme for the searches against databases of variable sizes.

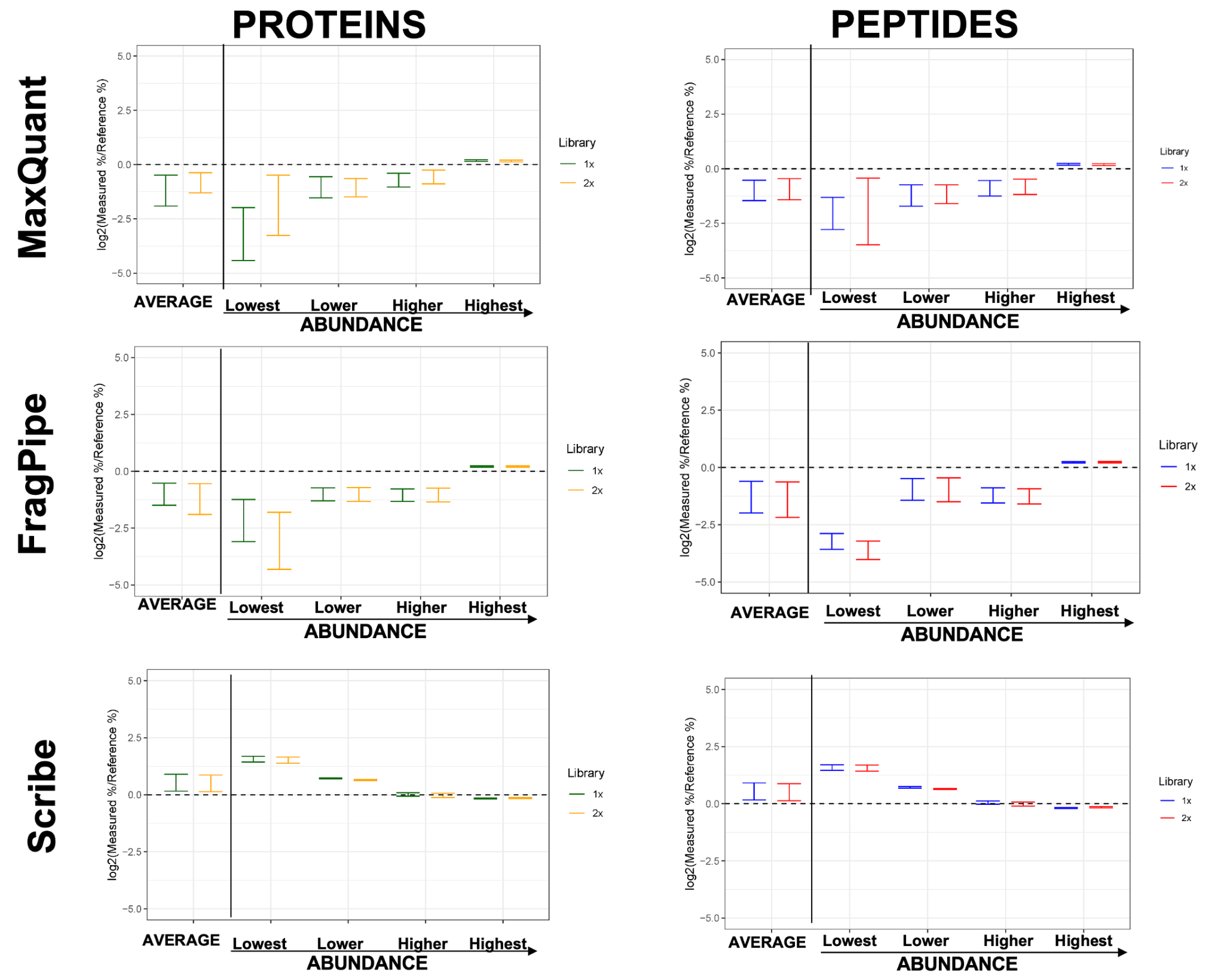

**Supplement S1 Figure S2. Quantitative analyses of the search results demonstrate a higher correlation between measured and reference values in the Scribe platform.** The protein-level and peptide-level intensities of the organisms of lowest abundance (Ne1, F0, ES18, P22, Nm1, BXL and Nu1); lower abundance (BS, NV, 841, PaD, DVH, Am2 and 137); higher abundance (KF7, CV, AK199, 259, HB2, VF, and PD) and highest abundance (CRH, ATN, K12, Pfl, SMS, Cup and LT2) were summed up. The percentage of the summed intensity at protein-level or peptide-level were compared to the expected percentage of the various tiers of organisms. The line at 0.0 indicates the absence of any difference between expected and measured values.
